# Morphometrical brain markers of sex difference

**DOI:** 10.1101/2020.08.01.232165

**Authors:** Daniel Brennan, Tingting Wu, Jin Fan

## Abstract

Many major neuropsychiatric pathologies, some of which appear in adolescence, show differentiated prevalence, onset, and symptomatology across the biological sexes. Therefore, mapping differences in brain structure between males and females during this critical developmental period may provide information about the neural mechanisms underlying the dimorphism of these pathologies. Utilizing a large dataset collected through the Adolescent Brain Cognitive Development study, we investigated the differences of adolescent (9-10 years old) male and female brains (*n* = 8325) by using a linear Support-Vector Machine Classifier to predict sex based on morphometry and image intensity values of structural brain imaging data. The classifier correctly classified the sex of 86% individuals with the insula, the precentral and postcentral gyri, and the pericallosal sulcus as the most discernable features. The role of these significant dimorphic features in psychopathology was explored by testing them as mediators between sex and clinical symptomology. The results demonstrate the existence of morphometrical brain markers of sex difference.

**Significance Statement:** Many psychiatric pathologies express differently across the sexes. Therefore, an understanding of the differences in brain structure between males and females during the critical developmental period of adolescence may provide the insights about the dimorphism of clinical symptomology and the general functions of the dimorphic brain structures. Using machine learning, we successfully classified males and females with a high accuracy based on morphometry and image intensity data extracted from structural MRI scans. The features which significantly contributed to classification were examined to determine brain regions which are dimorphic during adolescence. The relevance of these brain regions to the expression of psychopathology symptoms was also explored.

## Introduction

Many major neuropsychiatric pathologies are differentiated in prevalence, developmental trajectory and symptomatology across the biological sexes. For example, it is well documented that Autism Spectrum Disorder (ASD) shows a significantly higher prevalence in males compared to females (Werling & Geschwind, 2013). This imbalance has sparked various theories regarding the differences between male and female brains such as extreme male brain hypothesis (Baron-Cohen, 2002) and female protection effect (Robinson, Lichtenstein, Anckarsater, Happe, & Ronald, 2013), among others. In contrast, Major Depressive Disorder (MDD) shows an inverse pattern, with an approximate 2:1 ratio for females over males in lifetime prevalence (Picco, Subramaniam, Abdin, Vaingankar, & Chong, 2017; Seedat et al., 2009). This disparity is observed as early as adolescence (Salk, Hyde, & Abramson, 2017) and depressive symptoms may differ across the sexes (Martin, Neighbors, & Griffith, 2013). It is therefore immediately relevant to investigate sex differences in the brain. Determining whether brain markers for biological sex exist in adolescence is a crucial step for understanding differences in brain development between males and females. Establishing a pattern of brain dimorphism will point us towards possible sources of these divergent profiles of psychopathology and will lead to further understanding of about the functions of the dimorphic brain structures.

Univariate analyses have demonstrated consistent global morphological differences across men and women with disagreement in the magnitude and direction of localized differences in gray matter qualities (Kaczkurkin, Raznahan, & Satterthwaite, 2019; Lotze et al., 2019; Ruigrok et al., 2014), leading some studies to conclude that the overwhelming majority of the brain’s “mosaic” is largely overlapping between the sexes and localized regions of the brain are small and ultimately inconsequential (Joel et al., 2015). However, recent advances using machine learning multivariate classification methods demonstrate that indeed males and females do have differentiable brain features (Chekroud, Ward, Rosenberg, & Holmes, 2016), with classification accuracy exceeding 90% using structural brain imaging (Anderson et al., 2019; Chekroud et al., 2016). These studies have focused on samples with broad age ranges, including both pre and post-adolescent brain scans. While age is statistically accounted for in these models, established nonlinearities in brain development between males and females, e.g. (Gennatas et al., 2017; Kaczkurkin et al., 2019), do not permit a singular profile of dimorphism which is applicable across the lifespan. Instead, more focused investigation into brain dimorphism at specific timepoints during critical developmental periods, such as adolescence, are warranted to answer age-specific questions regarding brain dimorphism across biological sex.

In this study, we used a Support Vector Machine Classifier (SVC) to test the classification power of morphometrical features derived from structural T1 images of the ABCD sample. The Adolescent Brain Cognitive Development (ABCD) study provides a unique resource for characterizing the neural profile of the crucial development of adolescence (Hagler et al., 2019; Volkow et al., 2018). Brain morphometry and image intensity measures created via cortical surface reconstruction and subcortical segmentation using FreeSurfer’s automated pipeline (Fischl & Dale, 2000), which have been validated for use in children (Ghosh et al., 2010), were used as predictive features of biological sex. If the SVC could successfully classify biological sex significantly above chance using these features, this would demonstrate that dimorphism exists in the brain and there are measurable brain markers of sex during this critical developmental period. A mediation analysis was also conducted to explore the potential markers of biological sex that serve as mediators between sex and clinical symptoms measured by the Child Behavior Checklist (CBCL) syndrome scales and the Diagnostic and Statistical Manual of Mental Disorders, 5th Edition (DSM5)-oriented scales (see **Supplementary Methods** for details).

## Results

### SVC performance

Classification performance with all features and with each of the feature sets is shown in **Table 1**. An accuracy (mean ± standard deviation (SD)) of 86.3 ± 0.8% was obtained across the simulation of the full sample with all input features which was consisted of seven feature sets: cortical thickness, sulcal depth, cortical area, cortical volume, T1 grey matter intensity, T1 white matter intensity, and T1 gray matter and white matter contrast (GWC). Deep-learning did not improve performance compared to linear SVC: a training accuracy of 100% and test accuracy of approximately 85% were observed. Inspection of the precision and recall revealed no evidence of bias across sex in classification accuracy. Classification accuracy for all features was mostly retained after partitioning groups by clinical severity, and the linear SVC of the individual clinical tertials (low, middle, and high reported internalizing and externalizing behaviors) revealed similar classification accuracy across these groups (low = 82.9 ± 1.4%, med = 83.3 ± 1.4%, high = 82.4 ± 1.5%). Therefore, the examination of the features which significantly contributed to classification was based on the full sample rather than the stratified samples. All of the seven feature sets predicted significantly above chance level with a range of accuracy from 69.4% (white matter intensity) to 76.0% (cortical area).

**Table 1.**
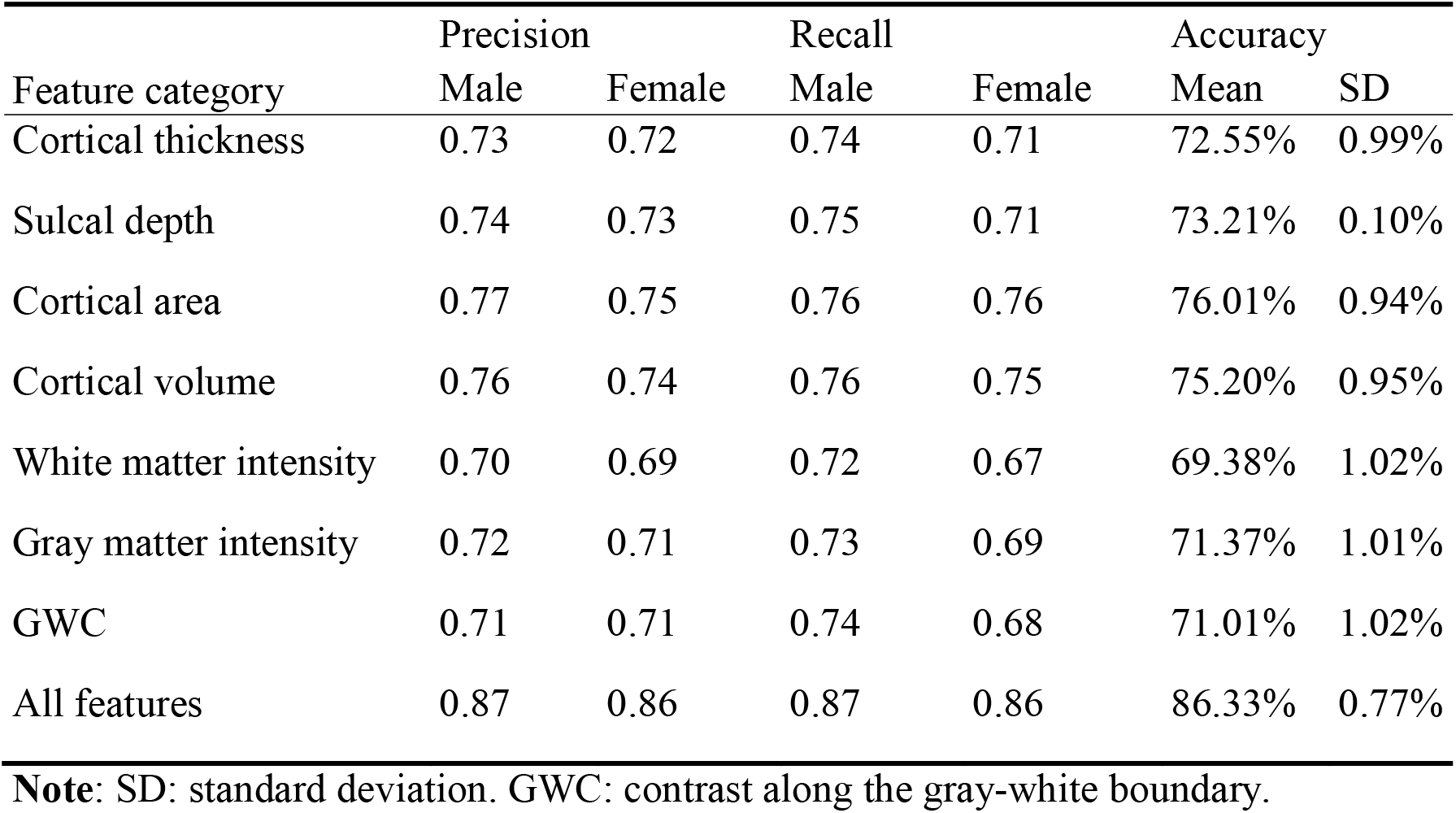
Support vector machine (SVM) classification performance.

| Feature category | Precision |  | Recall |  | Accuracy |  |
| --- | --- | --- | --- | --- | --- | --- |
|  | Male | Female | Male | Female | Mean | SD |
| Cortical thickness | 0.73 | 0.72 | 0.74 | 0.71 | 72.55% | 0.99% |
| Sulcal depth | 0.74 | 0.73 | 0.75 | 0.71 | 73.21% | 0.10% |
| Cortical area | 0.77 | 0.75 | 0.76 | 0.76 | 76.01% | 0.94% |
| Cortical volume | 0.76 | 0.74 | 0.76 | 0.75 | 75.20% | 0.95% |
| White matter intensity | 0.70 | 0.69 | 0.72 | 0.67 | 69.38% | 1.02% |
| Gray matter intensity | 0.72 | 0.71 | 0.73 | 0.69 | 71.37% | 1.01% |
| GWC | 0.71 | 0.71 | 0.74 | 0.68 | 71.01% | 1.02% |
| All features | 0.87 | 0.86 | 0.87 | 0.86 | 86.33% | 0.77% |

**Note:** SD: standard deviation. GWC: contrast along the gray-white boundary.

### Features that significantly contributed to classification

Four features with significant positive weights (female > male, regions in red) and nine features with significant negative weights (male > female, regions in blue) were identified and are visualized in **Figure 1** via the fsbrain visualization package (Schäfer, 2020). Regions with positive weights included the right postcentral gyrus (cortical thickness), the left long insular gyrus and central sulcus of the insula (cortical volume), the left anterior segment of the circular sulcus of the insula (cortical volume), and the temporal pole (GWC). Regions with negative weights included the left superior occipital gyrus (cortical thickness), the right lingual gyrus (sulcal depth), the left posterior ramus of the lateral sulcus (cortical area), the left precentral gyrus (cortical volume), the right pericallosal sulcus (gray matter intensity and white matter intensity), the left superior segment of the circular sulcus of the insula (GWC), the right lateral occipito-temporal gyrus (GWC), and the right superior segment of the circular sulcus of the insula (GWC). Labels and weights of these features are presented in **Table 2**. Mean feature weights across sex are visualized in **Figure 2** in order to demonstrate the difference in sensitivity between the machine-learning multivariate approach and univariate statistics: although these 13 features contributed to successful classification, not all variables are significantly different across groups. Some differences in feature means across groups would be detectable with a univariate analysis; however, many are largely overlapping or in unexpected directions, which would not be detectable using a univariate approach. The SVC is sensitive to non-linear dependencies between features, and is therefore able to capture more complex dimorphism within the brain. These features alone carry significant classification power with an average accuracy of 70.8 ± 1.0% observed in a linear SVC using only these 13 features as predictors of biological sex.

**Figure 1.**
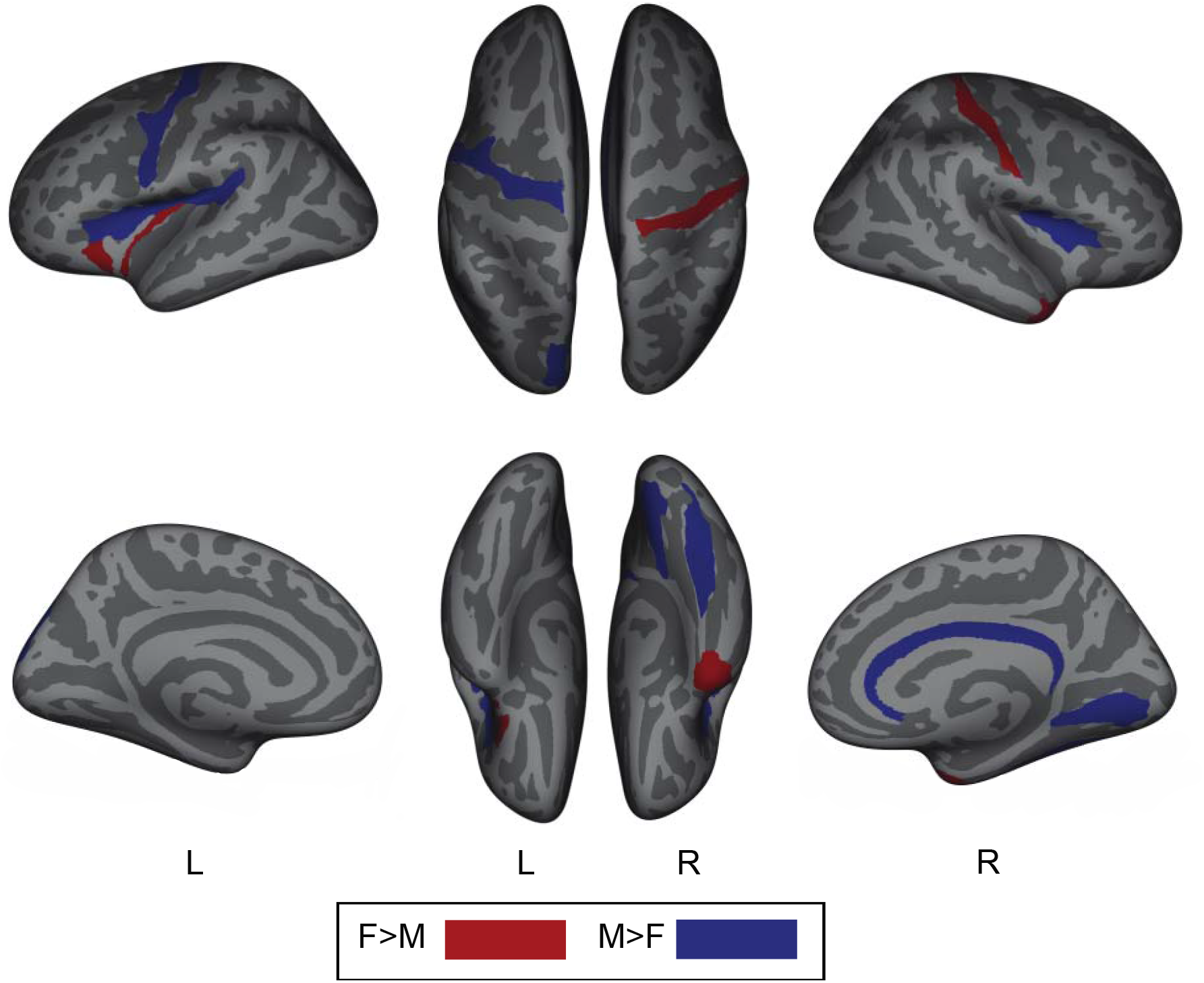
Dimorphic brain regions identified by SVM. Visualization of all brain regions with significant weights leading to successful support vector classification (SVC), independent of feature type. Red color signifies positive weight and therefore positive class contribution (female > male). Blue color signifies negative weight and therefore negative class contribution (male > female).

**Figure 2.**
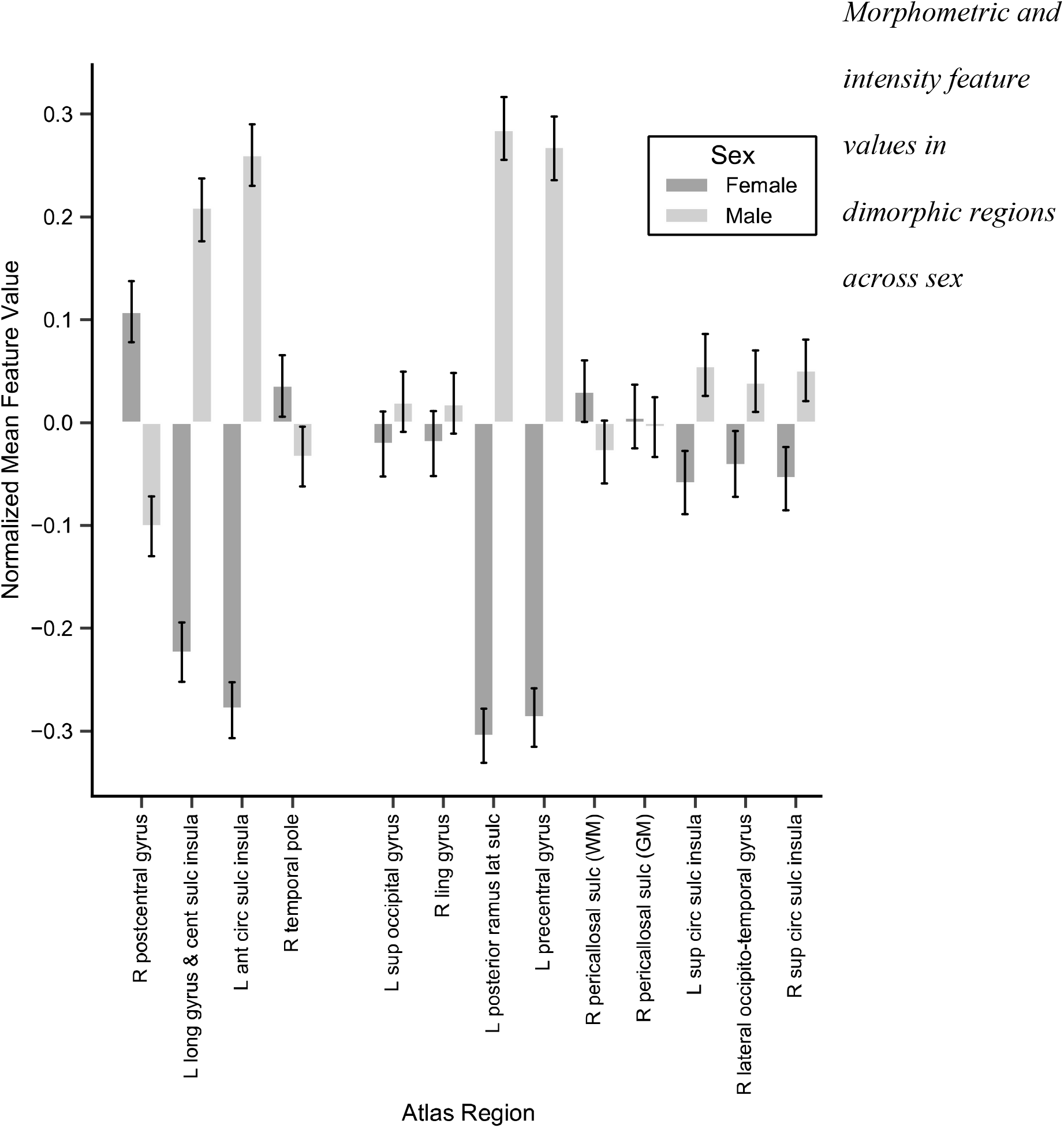
Morphometric and intensity feature values in dimorphic regions across sex. Univariate descriptive statistics (means and standard errors) of the 13 features identified in the SVC across sex.

**Table 2.**
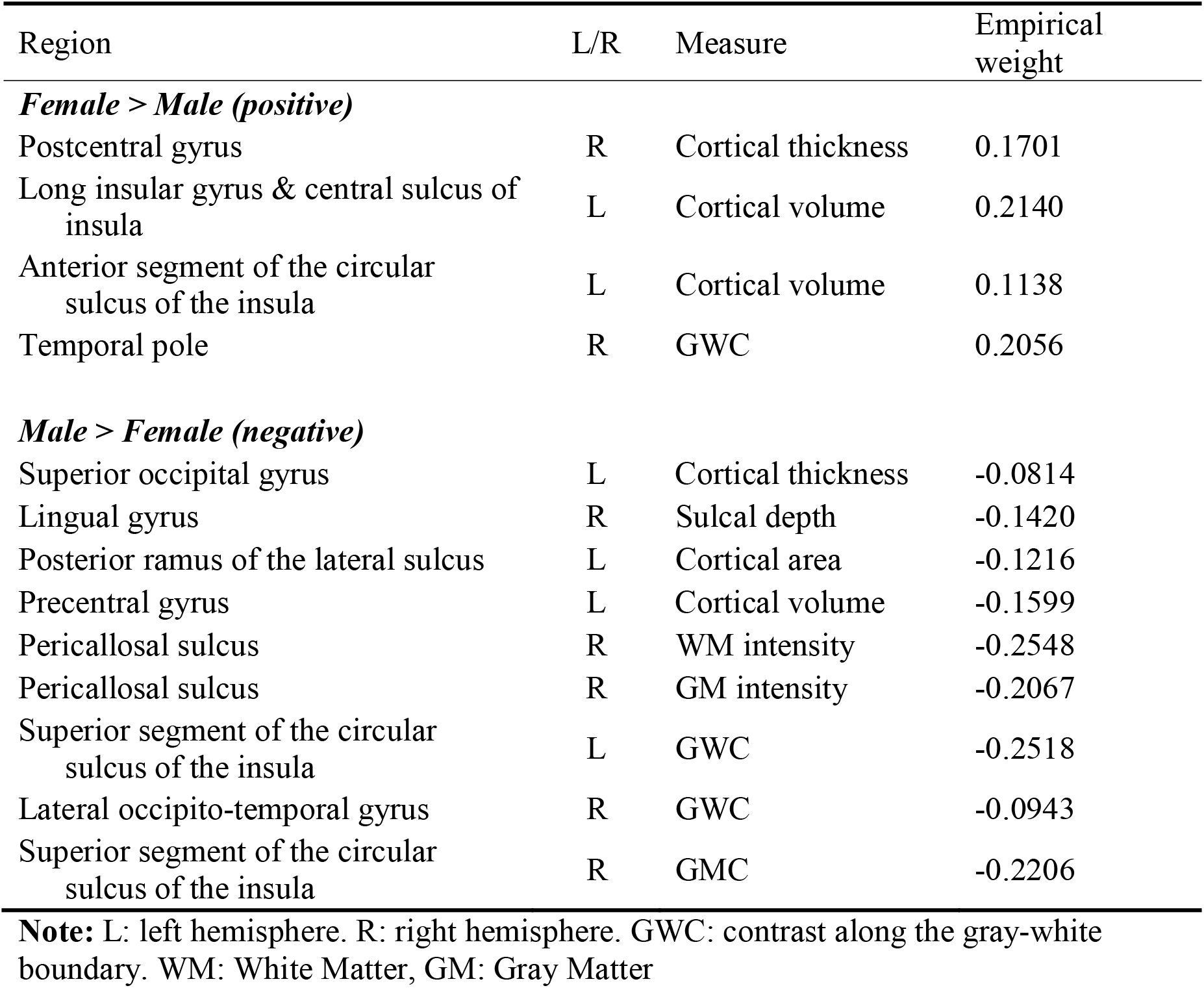
Weights significantly contributed to the SVC of males and females.

### Dimorphic features mediating the relationship between sex and clinical symptoms

The 13 regions identified as dimorphic were tested in parallel as potential mediators of the relationship between sex and clinical symptomology. The schema of the mediation analysis of the path from sex to brain morphometry and then to clinical symptomology is visualized in **Figure 3a**. This analysis revealed 3 unique significant morphometry features that served as the mediators including: GWC of the left superior segment of the circular sulcus of the insula on Child Behavior Checklist (CBCL) syndrome scales of withdrawn depressed and thought problems, CBCL total problems, and DSM-5-oriented scales of depression problems and stress (**Figure 3b**); surface area of the left posterior ramus of the lateral sulcus on CBCL syndrome scale of rule-breaking behavior and DSM-5-oriented scale of conduct problems (**Figure 3c**); and cortical volume of the left precentral gyrus and GWC of the left superior segment of circular sulcus of the insula on DSM-5-oriented scale of oppositional defiant problems (**Figure 3d**). See **Table 3** for details of the coefficients and *p* values.

**Figure 3.**
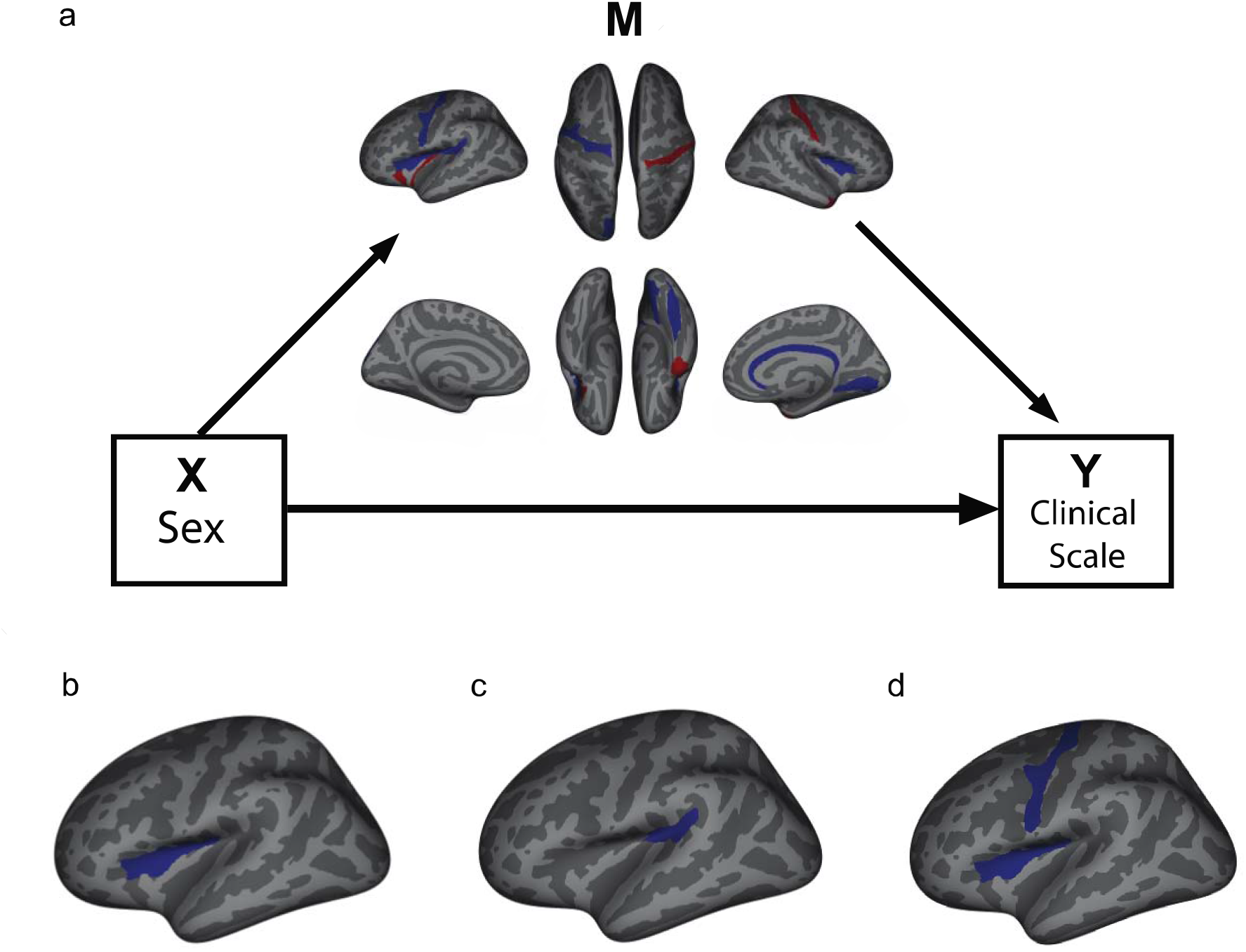
Mediation analysis schema and results. (a) Diagram of the multiple tested paths in the exploratory mediation analysis. The indirect path (Sex ⟶ Brain Feature ⟶ Clinical Scale) tests the parallel mediation effect of all dimorphic brain regions outlined our analysis on the Sex-Clinical Scale relationship, visualized in b, c, d. (b) GWC of the left superior segment of the circular sulcus of the insula on Child Behavior Checklist (CBCL) syndrome scales of withdrawn depressed and thought problems, CBCL total problems, and DSM-5-oriented scales of depression problems and stress; (c) surface area of the left posterior ramus of the lateral sulcus on CBCL syndrome scale of rule-breaking behavior and DSM-5-oriented scale of conduct problems; (d) and cortical volume of the left precentral gyrus and GWC of the left superior segment of circular sulcus of the insula on DSM-5-oriented scale of oppositional defiant problems.

**Table 3.** Results of exploratory mediation analysis for clinical symptomology.

| Y | M | Path | Coefficient | Uncorrected<br>*p* value |
| --- | --- | --- | --- | --- |
| CBCL Withdrawn Depression | Superior segment of the circular sulcus of the insula | c | -0.0015 | <0.001 |
|  |  | b | 25.6589 | 0.037 |
|  |  | a+b | -0.0363 | 0.040 |
| CBCL Thought | Superior segment of the circular sulcus of the insula | c | -0.0015 | <0.001 |
|  |  | b | 25.451 | 0.040 |
|  |  | a+b | -0.0375 | 0.024 |
| CBCL Rule breaking | left posterior ramus of the lateral sulcus | c | -90.4366 | <0.001 |
|  |  | b | -0.0007 | 0.083 |
|  |  | a+b | 0.0676 | 0.028 |
| CBCL Aggression | Superior segment of the circular sulcus of the insula | c | -0.0015 | <0.001 |
|  |  | b | 23.4542 | 0.037 |
|  |  | a+b | -0.0319 | 0.032 |
| CBCL Total Problems | Superior segment of the circular sulcus of the insula | c | -0.0015 | <0.001 |
|  |  | b | 50.9212 | 0.033 |
|  |  | a+b | -0.0731 | 0.032 |
| DSM Depression | Superior segment of the circular sulcus of the insula | c | -0.0015 | <0.001 |
|  |  | b | 26.9807 | 0.028 |
|  |  | a+b | -0.039 | 0.02 |
| DSM Oppositional Disorder | Left precentral gyrus | c | -576.8721 | <0.001 |
|  |  | b | -0.0001 | 0.171 |
|  |  | a+b | 0.0687 | 0.044 |
|  | Superior segment of the circular sulcus of the insula | c | -0.0015 | <0.001 |
|  |  | b | 26.8547 | 0.017 |
|  |  | a+b | -0.0364 | 0.02 |
| DSM Conduct Disorder | left posterior ramus of the lateral sulcus | c | -90.4366 | <0.001 |
|  |  | b | -0.0008 | 0.059 |
|  |  | a+b | 0.0831 | 0.016 |
| CBCL Stress | Superior segment of the circular sulcus of the insula | c | -0.0015 | <0.001 |
|  |  | b | 29.4708 | 0.019 |
|  |  | a+b | -0.042 | 0.012 |

**Note.** Path a: Sex on Brain region. Path b: Brain region on clinical. Path c: Sex on clinical, accounting for path a+b, e.g. total effect. Total mediation effect (indirect effect): path a+b

## Discussion

Using morphometric and image intensity values as predictors, we were able to classify biological sex with approximately 86% accuracy in an exceptionally large sample of 9-10 year-old males and females. We have also identified the 13 significant regional measures for the successful classification of males and females. As noted in prior findings relating to brain dimorphism (Anderson et al., 2019; Chekroud et al., 2016), local differences between the brains of males and females are not as inconsequential as hypothesized (Joel et al., 2015). We therefore conclude that there is sexual dimorphism in the human brain which is measurable during this critical developmental period. Contrary to previous analyses of biological sex and brain morphometry using machine learning (Anderson et al., 2019; Chekroud et al., 2016), the current study focuses exclusively on the developmental period of adolescence. This analysis therefore not only delineates brain markers of biological sex, but also contributes to the large body of research outlining unique trajectories of males and females during development. Our findings provide a whole-brain description of brain dimorphism which may be used to direct more pointed research into the relationship between brain dimorphism during adolescence and dimorphic clinical profiles which develop during this period.

There are multiple facets of the ABCD dataset which have contributed to the success of this analysis. An obvious advantage is the sheer number of individuals included in this study. At over 8,000 included observations, our analysis methods are well powered to reveal brain differences. Furthermore, the quantity and diversity of input features derived from the FreeSurfer pipeline are advantageous for two reasons. First, these brain measures can indicate independent features of brain morphometry, which can express a multitude of biological variables from a single scan type, e.g., T1 image. The inclusion of seven different T1 feature types undoubtedly increases the sensitivity of this analysis as demonstrated by this current study. On its own, any single feature type can only produce classification accuracy of as high as 76.01% while the predictive power of the aggregate of all feature types is clearly stronger. Second, the high dimensionality of the data afforded by the large diversity of features provides the benefit of more efficient linear separability while also not succumbing to the “curse of dimensionality” due to an adequate number of observations. This facet of the data structure is not to be overlooked. Through the possibility of linearly separating our data, we gain explainability within our model not afforded by other non-linear methods, e.g., kernelized SVC, while retaining the ability to detect nonlinear dependencies on classification based on these features with the high dimension SVC. Indeed, this advantage over univariate approaches is best understood by inspecting feature weights and feature means simultaneously. Positive or negative weights leading to female or male prediction do not necessarily have analogous differences of means. Attempting similar analyses utilizing the univariate method would therefore not be able to detect such differences. The “brain mosaic” is truly a complex, yet robustly observable, non-linear combination of features.

The relationship between sexual dimorphism in the brain and behavioral dimorphism is complex and not yet fully understood (De Vries, 2004; Velasco, Florido, Milad, & Andero, 2019; Yagi & Galea, 2019), although there appears to be a relationship between neuroendocrine expression during development and the limbic system which has implications for risk-taking (Casey, Jones, & Hare, 2008) and clinical symptoms, such as anxiety (Spielberg, Schwarz, & Matyi, 2019). Several cortical regions identified by our analysis as being dimorphic have direct relationship with the limbic system, including multiple sulci in the left insula and left lateral sulcus, the circular sulcus of the right insula, left pre-central sulcus, and right post-right central sulcus, and right peri-callosal sulcus. The limbic system is implicated in sexual function (George & Lorberbaum, 2002), and anatomical dimorphism within the limbic system in mammalian species is well documented (Cahill, 2006; Madeira & Lieberman, 1995). While sex-specific investigations exist between cortico-limbic circuitry and clinical symptoms during adolescence (Mohamed Ali, Vandermeer, Sheikh, Joanisse, & Hayden, 2019), it is still important to uncover evidence linking brain dimorphism present during adolescence to the development of clinical symptoms which are commonly observed or developed during this developmental period. Therefore, relating the regional measures identified as dimorphic within this sample to differences in clinical symptomology and behavioral clinical precursors is a necessary step in outlining which may underly the development of dimorphic clinical profiles across in adolescence and beyond. We observed several possible relationships between sex and clinical symptomology, mediated by the dimorphic brain regions uncovered by our classification analysis. The results of this analysis point towards two main regions with implications for dimorphic symptomology: posterior ramus of the lateral sulcus, and most prominently, the GWC within the insula. The necessity of the insula has been demonstrated through lesion studies showing significant disruption of addictive behaviors (Naqvi, Rudrauf, Damasio, & Bechara, 2007), cognitive control (Wu et al., 2019) and acquisition of taste aversion (Roman & Reilly, 2007). GWC is thought to reflect the differential myelination of the cerebral cortex and subjacent WM (Norbom et al., 2019), and individual differences in GWC within the insula, cingulate and pre/post central cortices are associated with both mental health and general cognitive functioning in adolescents (Norbom et al., 2019).

The insula is a multifaceted region responsible for a diversity of functions (Flynn, 1999): the insula is central to autonomic and visceral information processing and their integration, highlighted by its subcortical, limbic, and brain stem connections. Its cortical connections are predominantly with other neocortical areas. Insular-cortical and sub-cortical connections, especially with the thalamus and basal ganglia, underscore the posterior insula’s role in somatosensory, vestibular, and motor integration. The circular sulcus outlines the circumference of the insula, separating it from the temporal, frontal and parietal lobes. Indeed, the many structural connections of the insula underly its role as a hub for various specialized functions including attention, cognitive, affective and regulatory functions (Menon & Uddin, 2010). Recent evidence has placed the insula as an important component of emotional processing in the brain (Gu, Hof, Friston, & Fan, 2013). The insular cortex has also been shown to have a role in social processing (Spagna et al., 2018), predicting social emotions in others (Lamm & Singer, 2010) and is necessary for empathetic pain perception (Gu et al., 2012). The functional integration of cognitive, regulatory and affective functions within this region (Kurth, Zilles, Fox, Laird, & Eickhoff, 2010) supports a process known as interoceptive awareness (Gu, Liu, Van Dam, Hof, & Fan, 2013; Wang et al., 2019). It is posited that interoceptive awareness gives rise to various motivations and drives, as well as emotional and affective experiences through the maintenance of homeostatic balance within the body (Khalsa et al., 2018). The insula is therefore a possible source of disfunction in affective and emotional disorders which have close relationships with the body (Khalsa et al., 2018; Paulus & Stein, 2010). The roles of the insula presented here provide theoretical backing to the results of our exploratory analysis, which demonstrates a mediating role of GWC in the circular sulcus of the insula between sex and several dimensions of clinical symptomology. It is therefore relevant to carefully consider the individual differences within the insular cortex across sex and the possible role this region plays in the dimorphic development and profiles of psychopathology.

## Materials and Methods

### Demographics of the sample

The ABCD study is the largest project of its kind to investigate the neurobehavioral development of adolescents with multimodal brain imaging data collected from an exceptionally large sample of 9-10 year old children (n = 11875) (Casey et al., 2018). In addition, the ABCD dataset was created in part to identify early markers of substance abuse and other mental health problems (Barch et al., 2018; Lisdahl et al., 2018) with recruitment protocols reflecting the intention to include a large number of children who show early signs of externalizing and internalizing symptoms (Loeber et al., 2018). The ABCD dataset is a well-controlled and thorough resource, and therefore is optimal for analysis of sex differences during this critical age range using machine learning.

Parcellated structure MRI data of 11,533 participants were provided in the ABCD dataset release 2.0. We have applied the following exclusion criteria to the data: (1) participants with no T1 scan passing raw data quality control; (2) participants not passing the FreeSurfer quality control; (3) participants with missing data for T1 related morphometric or intensity features; (4) participants with missing data for cognitive and clinical measures. The final sample consisted data from 8325 participants (mean ± SD) age = 119.07 ± 7.48 months), including 4308 males (age = 119.15 ± 7.46 months) and 4017 females (age = 118.99 ± 7.50 months). The difference in age (a strong influence on brain morphometry) between male and female groups was not significant, *t*(8325) = 0.95, *p* = 0.34, with a Bayes Factor value of 0.04. Demographic information for the included participants was reported in **Supplementary Table 1**.

### MRI data of the sample

Fully preprocessed morphometric and image intensity values of the structural T1 images were provided by the ABCD study (Hagler et al., 2019), and were included as the predictive features in this analysis (see **Supplementary Matierials** for MRI acquisition details). Morphometric and image intensity measures were created from structural T1 images via cortical surface reconstruction using FreeSurfer’s automated pipeline (Fischl & Dale, 2000). Labels for cortical gray matter and underlying white matter voxels were assigned based on surface-based nonlinear registration to the Destrieux atlas (Destrieux, Fischl, Dale, & Halgren, 2010). Morphometric measures included four categories: cortical thickness, cortical surface area, cortical volume, and sulcal depth. While these measures are correlated, they indicate different facets of gray matter integrity, and some (surface area/thickness) are genetically independent (Winkler et al., 2010). Image intensity measures included three categories: average gray matter intensity, average white matter intensity, and the average contrast along the gray-white boundary: GWC = (white – gray)/(white + gray). Intensity measures were included because they provide independent information relative to morphometry and likely reflect local influences on morphometry (Westlye et al., 2009). Additionally, GWC is a sensitive measure of local variation in tissue integrity and myelin degradation (Uribe et al., 2018), which likely reflects a unique biological signal which varies significantly with age (Vidal-Pineiro et al., 2016) and may have increased clinical sensitivity relative to cortical thickness (Makowski et al., 2019). Each category of feature was provided for each of the 148 regions of interest (ROIs, 74 ROIs for each hemisphere). An average across ROIs in the left hemisphere, an average across ROIs in the right hemisphere, and an average across all ROIs for each morphometric measure were also included as the predicting features.

### Clinical Measures

The Achenbach System of Empirically Based Assessment (ASEBA) is one of the assessments of mental health in the ABCD dataset, which comprises the CBCL and the DSM-5-oriented scales (see **Supplementary Matierials** for more details). The CBCL is a parent-reported scale used with children 6 to 18 years old (CBCL/6-18), which is made up of eight empirically-based syndrome scales: anxious/depressed, withdrawn depressed, somatic complaints, thought problems, attention problems, rule-breaking behavior, and aggressive behavioral. Those scales group into three composite scales: internalizing (sums the first three scales), externalizing (sums the last two scales), and total problems (sums all scales). The DSM-5-oriented scales include nine diagnostic categories: depression problems, anxiety disorders, somatic problems, attention deficit/hyperactivity problems (ADHD), oppositional defiant problems, conduct problems, sluggish cognitive tempo, obsessive-compulsive problems, and stress. In our included sample, 74.76% of participants provided this clinical data within 6 months of scanning; 16.60% completed within 6-12 months, and 8.64% completed between 12-18 months. Both raw scores and the age normed t-scores of each of these measures were provided in the ABCD dataset.

### Classification with Support Vector Machine

SVC is a classification algorithm that attempts to find a hyperplane which best separates binary classes in hyperspace of predictive features. The above mentioned four categories of morphometry measures and three categories image intensity values derived from ABCD’s pipeline were used as predictive features for binary classification of sex within this sample. A total of 1057 features were standardized across all participants and entered into a linear SVC via the scikit-learn machine learning library (Pedregosa et al., 2011) in Python to predict the sex of each individual (parameter C = 0.01). The target variable sex was coded as “1” for females and “0” for males. Classification decisions in linear SVC are computed through a linear combination of weights and input data: y□ = *w*^*T*^x+b. If the result y□ is positive, the target class (female) is output as prediction; negative values of y□ lead to non-target (male) class prediction. Therefore, positive weights can be interpreted as contributing to target class decisions and negative weights can be interpreted as contributing to non-target class decisions. This facet of linear SVC calculation was utilized in a subsequent analysis to determine multivariate features which influenced the classification of both females and males, and therefore represent dimorphic brain regions. The performance of the SVC was assessed using a simulation method of 10,000 iterations using cross validation. That is, the classifier was trained on a random subset of the data (80% of observations) and tested on the remaining subset (20% of observations) for each independent iteration of the simulation. Classification metrics (accuracy, precision, recall) and calculated feature weights for each iteration were recorded across this simulation.

Two additional analyses were conducted to verify the above linear SVC approach. First, linear SVC performance was compared to deep-learning to examine whether improved classification performance can be achieved through non-linear machine learning. All features were included. A fully connected dense network created using TensorFlow 2 (Abadi et al., 2016) (2 hidden layers, each size 128, Relu activation, output = size 2, softmax activation; loss = sparse categorical cross-entropy, optimized with Adam, batch normalized) was trained and tested on an identical split (80% train, 20% test). Second, as children who exhibit internalizing and externalizing symptoms were specifically recruited for this sample (Loeber et al., 2018), these demographics are overrepresented and may influence classification performance. To ensure no bias in classification due to differences in internalizing and externalizing severity, the linear SVC simulation procedure with all features included was reproduced on three tertiles of the original dataset: low, middle and high levels of reported internalizing and externalizing behaviors (internalizing + externalizing raw score; Low: 0-3, Medium: 3-10, High: 10-83). Lastly, we identically applied the above SVC procedure to each of the seven input feature sets to evaluate the individual importance and contribution of these different feature types.

### Determination of significant contributing features

An advantage of linear SVC is that non-zero feature weights may be interpreted as contributing to successful classification (Géron, 2017). A second simulation was conducted using a permutation method to assess the importance of observed feature weights for successful classification: mean observed (empirical) feature weights from the first simulation (averaged across the 10,000 iterations) were compared to null distributions of feature weights. These null distributions of feature weights were created by replicating the SVC procedure using random assignment of classification labels (sex) over a second simulation of 10,000 iterations. Mean empirical weights outside a 95% confidence interval for each feature (Bonferroni corrected: *n* = 1057; mean ± 4.069 SD) determined by the null distribution of sample weights were considered to be significant contributors to successful classification.

### Clinical implications of sexually-dimorphic brain features

Exploring dimorphism in the brain may lead to insights regarding dimorphism within well documented differences in clinical symptomology and development across sexes. Using the brain features uncovered by the linear SVC, we performed an exploratory investigation assessing the role of these dimorphic brain regions in mediating the relationship between sex and clinical symptomology. For each of the clinical measures, we performed a mediation analysis with all significant dimorphic features identified by linear SVC as potential parallel mediators (M) for the effect of biological sex (X) on the clinical measure (Y). Significant indirect effects (X⟶M⟶Y) were computed using a bias-corrected, non-parametric bootstrap method (Hayes & Rockwood, 2017) via the Pingouin statistical package (Vallet, 2018). Significant indirect effects were reported regardless of significance of direct or total effect (Loeys, Moerkerke, & Vansteelandt, 2014). A liberal threshold (*p* < 0.05, uncorrected) was adopted for this exploratory analysis as we aimed to demonstrate the potential role of dimorphic regions in clinical symptomology.

## Supplementary Information

### Clinical questionnaires

Clinical symptomology was assessed using the Achenbach System of Empirically Based Assessment (ASEBA), which comprise the Child Behavior Checklist (CBCL) and DSM-5-oriented scales (DSM-5) (T. M. Achenbach, Dumenci, & Rescorla, 2003). The CBCL (Thomas M. Achenbach, 1991; Thomas M. Achenbach & Rescorla, 2001) is a parent-completed rating scale of behavior which qualifies internalizing and externalizing behaviors. Though empirically derived, these scales have weak relationship to the diagnostic standards of the DSM, and thus the DSM-5 oriented scales were created (Ebesutani et al., 2010).

### MRI acquisition parameters

Structural MRI data were downloaded from ABCD Data Release 2.0 (3/26/2019) which were collected from participants of the ABCD study (for the outline, see Casey et al. (2018)). To increase scanner completion, participants completed pre-scan training in a mock-scanner. During scan-time, a child friendly movie was turned on as the child entered the scanner and remained on during acquisition of the localizer and 3D T1 scan. For the structural scan, 3D T1-weighted magnetization-prepared rapid acquisition gradient echo scan was obtained (176 slices at 256 × 256 mm field of view; 1.0 mm isotropic, flip angle = 8.0°. Total acquisition time = 7 min and 12 sec).

## Supplementary Tables

**Supplementary Table 1.** Demographics of Adolescent Brain Cognitive Development (ABCD) sample.

| Race | Gender |  |  |
| --- | --- | --- | --- |
|  | Male | Female | Male + Female |
| White | 3348 (38.82%) | 3026 (35.08%) | 6374 (73.90%) |
| Black/African American | 948 (11.00%) | 909 (10.54%) | 1857 (21.53%) |
| American Indian | 145 (1.68%) | 143 (1.66%) | 288 (3.34%) |
| Alaska Native | 4 (0.05%) | 1 (0.01%) | 5 (0.06%) |
| Native Hawaiian | 1 (0.01%) | 7 (0.08%) | 8 (0.10%) |
| Guamanian | 4 (0.05%) | 0 (0) | 4 (0.05%) |
| Samoan | 15 (0.17%) | 2 (0.02%) | 17 (0.20%) |
| Pacific Islander | 51 (0.60%) | 14 (0.16%) | 65 (0.75%) |
| Asian Indian | 79 (0.92%) | 37 (0.43%) | 116 (1.34%) |
| Chinese | 56 (0.65%) | 74 (0.86%) | 130 (1.51%) |
| Filipino | 56 (0.65%) | 51 (0.59%) | 107 (1.24%) |
| Japanese | 27 (0.31%) | 32 (0.37%) | 59 (0.68%) |
| Korean | 35 (0.41%) | 39 (0.45%) | 74 (0.86%) |
| Vietnamese | 20 (0.23%) | 28 (0.32%) | 48 (0.56%) |
| Asian: Other | 39 (0.45%) | 32 (0.37%) | 71 (0.82%) |
| Other | 306 (3.55%) | 288 (3.34%) | 594 (6.89%) |
| Refuse to Answer | 25 (0.29%) | 19 (0.22%) | 44 (0.51%) |
| Don't Know | 30 (0.35%) | 36 (0.42%) | 66 (0.77%) |

**Note:** Demographics of ABCD sample based on parent questionnaire. Columns will not summate to total subject number, because responses for a child's race are not mutually exclusive. Percentages may therefore be interpreted as the percentage of sample which is identified by parent as a given race.

## Acknowledgements

Data used in the preparation of this article were obtained from the Adolescent Brain Cognitive Development (ABCD) Study (https://abcdstudy.org), held in the NIMH Data Archive (NDA). A listing of participating sites and a complete listing of the study investigators can be found at https://abcdstudy.org/scientists/workgroups/. ABCD consortium investigators designed and implemented the study and/or provided data but did not necessarily participate in analysis or writing of this report. This manuscript reflects the views of the authors and may not reflect the opinions or views of the NIH or ABCD consortium investigators.

